# Internal noise in contrast discrimination propagates forwards from early visual cortex

**DOI:** 10.1101/364612

**Authors:** Greta Vilidaite, Emma Marsh, Daniel H. Baker

**Author notes:** **Corresponding author** Daniel H Baker.

## Abstract

Human contrast discrimination performance is limited by transduction nonlinearities and variability of the neural representation (noise). Whereas the nonlinearities have been well-characterised, there is less agreement about the specifics of internal noise. Psychophysical models assume that it impacts late in sensory processing, whereas neuroimaging and intracranial electrophysiology studies suggest that the noise is much earlier. We investigated whether perceptually-relevant internal noise arises in early visual areas or later decision making areas. We recorded EEG and MEG during a two-interval-forced-choice contrast discrimination task and used multivariate pattern analysis to decode target/non-target and selected/non-selected intervals from evoked responses. We found that perceptual decisions could be decoded from both EEG and MEG signals, even when the stimuli in both intervals were physically identical. Above-chance decision classification started <100ms after stimulus onset, suggesting that neural noise affects sensory signals early in the visual pathway. Classification accuracy increased over time, peaking at >500ms. Applying multivariate analysis to separate anatomically-defined brain regions in MEG source space, we found that occipital regions were informative early on but then information spreads forwards across parietal and frontal regions. This is consistent with neural noise affecting sensory processing at multiple stages of perceptual decision making. We suggest how early sensory noise might be resolved with Birdsall’s linearisation, in which a dominant noise source obscures subsequent nonlinearities, to allow the visual system to preserve the wide dynamic range of early areas whilst still benefitting from contrast-invariance at later stages. A preprint of this work is available at: http://dx.doi.org/10.1101/364612

## 1 Introduction

The ability to make comparisons between sensory stimuli of different intensities has profound survival value for most organisms. Animals might benefit from choosing the ripest fruit based on colour, swimming towards the warmest patch of ocean, or selecting the mate with the loudest roar. Understanding the features of the central nervous system that limit such sensory discriminations has been a focus of research in many areas of psychology and neuroscience, from early work in humans (c.f. Weber’s law, Fechner, 1912), and experiments with model organisms (Busse et al., 2011; Hecht & Wald, 1934) to studies using contemporary neuroimaging techniques (Boynton, Demb, Glover, & Heeger, 1999).

A widely studied perceptual task is the ability to discriminate between visual stimuli of different contrasts. Human contrast discrimination performance is constrained by the nonlinearity that maps physical contrast to neural response, and the intrinsic variability of the neural representation (‘internal noise’). Psychophysical, neurophysiological and neuroimaging work have converged on a nonlinearity that is expansive at low contrasts and compressive at higher contrasts (Boynton et al., 1999; Busse, Wade, & Carandini, 2009; Legge & Foley, 1980). However, there is substantially less agreement regarding the details of performance-limiting internal noise.

Most psychophysical models make the assumption that the dominant source of noise for contrast discrimination is additive (i.e. independent of signal strength) and impacts at a late stage of processing. The primary justification for this arrangement is the observation that a dominant source of noise occurring before a nonlinearity will neutralise the effects of that nonlinearity, rendering it invisible to inspection (termed Birdsall’s theorem; Klein & Levi, 2009; Smith & Swift, 1985). Since contrast transduction is observably nonlinear (Boynton et al., 1999; Busse et al., 2009; Legge & Foley, 1980), any early sources of noise must be negligible in comparison to the magnitude of late additive noise.

On the other hand, most electrophysiological and neuroimaging studies have suggested that perceptually relevant noise is located in early sensory areas (Campbell & Kulikowski, 1972; Carandini, 2004; Roelfsema & Spekreijse, 2001). Ress and Heeger (2003) demonstrated the influence of early sensory noise by measuring fMRI blood-oxygen-level dependent (BOLD) responses in areas V1-V4 during contrast detection. They found that *false alarms* (trials on which the stimulus was absent, but reported as seen) evoked higher responses than *misses*, (trials on which the stimulus was present, but reported as not seen) suggesting that these areas encoded perceptual experience of the stimuli rather than the presence of the stimulus itself. The origin of the spurious activity in the case of *false alarms* is presumably neural noise in these early areas. Similarly, several intracranial primate electrophysiology studies have been able to predict the perceptual decisions of monkeys from neural activity recorded in early visual areas (Britten, Newsome, Shadlen, Celebrini, & Movshon, 1996; Britten, Shadlen, Newsome, & Movshon, 1992; Michelson, Pillow, & Seidemann, 2017). This suggests that sensory decisions are influenced by neural noise at an early stage of processing.

In this study, we attempt to understand how neural activity governs observer responses in a two-interval-forced-choice (2IFC) contrast discrimination paradigm, using methods typical of such studies. Two stimuli are presented in a random order, one containing a ‘pedestal’ of a fixed contrast (here 50%), and the other containing the pedestal plus a ‘target’ contrast increment. This paradigm involves several complicating factors that must be considered, including: (i) the observer must retain a neural representation of the first stimulus for comparison with the second stimulus, (ii) individuals might have idiosyncratic biases to prefer one or other interval, and (iii) fast acting adaptation (often termed repetition suppression) effects might reduce the neural response to the second stimulus (and perhaps also its appearance). We recorded evoked responses using both EEG (Experiment 1) and MEG (Experiment 2). We perform traditional univariate analyses, and also employ multivariate pattern analysis to decode participants’ percepts. Advantages of pattern analysis are that it can detect subtle and complex effects that might be missed by univariate analyses, is expressed in standard units (classifier decoding accuracy) that are independent of imaging modality, and permits testing of pattern generalisation across conditions and time (King & Dehaene, 2014). The high temporal resolution (∼1ms) of electromagnetic recording techniques enabled us to closely examine the timecourse of perceptual decision making, and the spatial resolution of MEG source space allowed us to investigate the involvement of discrete anatomical brain areas.

Our primary motivation was to determine whether the dominant source of neural noise is located in early sensory brain areas, or later (more frontal) areas involved in making decisions. To achieve this, our most crucial experimental condition is one in which the target contrast increment is 0%, meaning that the two stimuli to be compared contain only the pedestal and are therefore physically identical. Any differences in the neural representation that correspond to perceptual decisions must be due to processes occurring within the participant’s nervous system, rather than due to differences in the stimulus. We also included conditions in which the target contrast was >0% in order to measure psychophysical accuracy, to keep participants motivated, and to provide information on the timecourse of contrast discrimination when physical stimuli differ.

## 2 Methods

### 2.1 Participants

Twenty-two adults with normal or corrected-to-normal vision took part in Experiment 1 and ten took part in Experiment 2. All participants gave written informed consent. Experiment 1 was approved by the Ethics Committee of the Department of Psychology at the University of York, and Experiment 2 was approved by the York Neuroimaging Centre Ethics Committee.

### 2.2 Stimuli and psychophysical task

Stimuli were horizontally oriented sine wave gratings with a spatial frequency of 1c/deg and a diameter of 10 degrees. The edges of the gratings were blurred by a cosine function. On each trial, two stimuli were presented: a pedestal stimulus of 50% contrast (where percent contrast is defined as 100*(*L*_*max*_–*L*_*min*_)/(*L*_*max*_+*L*_*min*_), where *L* is luminance), and a pedestal+target stimulus consisting of the 50% contrast pedestal plus a target contrast increment. Five target contrast conditions were used in Experiment 1: 0% (no target), 2%, 4%, 8% and 16%. In Experiment 2 only the 0% (no target) and 16% target contrast conditions were used. Note that in the ‘no target’ conditions, the stimuli displayed were physically identical and the ‘target’ interval assignment was arbitrary. Participants were not informed of this, and still made a judgement about which interval appeared higher in contrast.

The two stimuli on each trial were presented sequentially for 100ms each, with a random inter-stimulus interval between 400ms and 600ms. The inter-trial interval followed the participant’s response, and was of variable length between 1000ms and 1200ms to avoid distortion of ERP averages (Woldorff, 1993). The order of target and non-target intervals within trials was counterbalanced. Trials of different target contrasts were intermixed and the order was randomized. Stimulus onsets and participant responses were recorded on the M/EEG trace using low-latency digital triggers.

### 2.3 EEG data collection

Event-related potentials were recorded using an ANT Neuroscan EEG system and a 64-channel Waveguard cap with electrodes arranged according to the 10/20 system. The ground electrode was positioned at *AFz*, and a whole head average was used as a reference. Data were digitised at 1kHz using the *ASALab* software. Stimuli were presented on a ViewPixx 3D display (VPixx Technologies Inc., Quebec, Canada) running in M16 mode (16-bit luminance resolution) with a mean luminance of 51cd/m^2^ and a refresh rate of 120Hz, using Matlab and elements of the *Psychophysics Toolbox* (Brainard, 1997; Kleiner, Brainard, & Pelli, 2007; Pelli, 1997). The display was gamma corrected using a Minolta LS110 photometer, fitting the data with a 4-parameter exponential function, and transforming stimulus intensities using the inverse of the function to ensure linearity.

Participants were seated in a darkened room 57cm away from the display. Instructions for the task were to ‘indicate the grating that appeared higher in contrast’. They were asked to fixate on a central cross throughout the task and used a mouse to indicate their responses. There were 200 trials per target contrast (1000 trials total, yielding 2000 stimulus-locked ERPs). The task was run in 5 blocks of approximately 8 minutes, with short breaks in between.

### 2.4 MEG data collection

MEG data were recorded using a 4D Neuroimaging Magnes 3600 Whole Head 248 Channel MEG scanner housed in a purpose-built Faraday cage. The data were recorded at 1017.25Hz, with 400Hz Bandwith using a High Pass DC filter. Nine channels were identified as having failed and were removed from all analyses. The location of the head inside the dewar was continuously monitored throughout the experiment using 5 position indicator head coils. Stimuli were presented on an Epson EB-G5900 3LCD projector (refresh rate 60Hz; mean luminance 160cd/m^2^) with a 2-stop ND filter, using *Psychopy* v1.84 (Peirce, 2007). The projector was gamma corrected using a Minolta LS110 photometer, fitting the data from each channel (red, green and blue) with a separate exponential function, and transforming stimulus intensities using the inverse of the function to ensure linearity.

Participants were seated in a hydraulic chair in front of the projector screen in a dark room. Prior to the task the three dimensional shape of the participant’s head was registered using a Polhemus Fasttrack headshape digitization system. Five fiducial points were used for this over two registration rounds. If the distance in location between the first and second round was >2mm, the registration was repeated. When successful, the headshape was then traced and recorded using a digital wand. This was later coregistered with T1-weighted anatomical MRI scans of each participant acquired in separate sessions using a 3T GE Signa Excite HDx scanner (GE Healthcare).

Participants fixated on a small central cross throughout the task. The experiment was completed in a single block consisting of 240 trials per contrast condition (480 trials in total, yielding 960 stimulus-locked ERPs), with a total acquisition duration of around 20 minutes. A single hand response pad was used to make responses in the experiment.

### 2.5 EEG data analysis

EEG recordings were bandpass filtered from 0.01Hz (cosine ramp) to 30Hz (Hanning window). They were then epoched into 1 second-long windows (200ms before stimulus onset to 800ms after) for each interval of every trial. Each epoch was then baselined at each electrode independently by subtracting the mean response over the 200ms preceding stimulus onset. ERPs were then sorted by target/non-target intervals for stimulus classification analysis and then again by selected/non-selected intervals for decision classification. No artifact rejection was performed, as we have generally found in previous studies (e.g. Coggan, Baker & Andrews, 2016) that this has no material impact on classification accuracy when trial numbers are large, stimulus presentations are brief, and participants are adults (as here).

To perform univariate analyses, ERPs were averaged across a cluster of 10 posterior electrodes (Oz, O1, O2, POz, PO3-8), and significance was determined using cluster corrected paired-samples t-tests across participants (Maris & Oostenveld, 2007). The significance of each cluster was determined by comparing to a null distribution of summed t-values derived by randomly permuting the labels of the largest cluster 1000 times. To perform multivariate analyses, a support vector machine (SVM) was used to classify the data independently at each sample point (i.e. in 1ms steps). A second stage of normalization was applied at each time-point and each electrode by subtracting the mean response across all intervals and conditions for that time/sensor combination. The data were then randomly averaged in five subsets of 40 trials for each category (target/non-target or selected/non-selected), of which four subsets were used to train the model and one was used to test it. The classifier algorithm creates a parameter space of all data points and then fits a hyperplane boundary that maximizes the distances between the support vectors of each category. Classifier accuracy for categorising the test data was averaged across 1000 repetitions of this analysis (with different random allocations of trials on each repetition), and was repeated for each target contrast condition. We used the same non-parametric cluster correction procedure as for the univariate analyses (Maris & Oostenveld, 2007) to identify time periods where classifier accuracy was significantly above chance (using t-tests across participants). We then averaged timecourses across participants for visualization purposes.

### 2.6 MEG data analysis

Cortical reconstruction and volumetric segmentation was performed with the *Freesurfer* image analysis suite (http://surfer.nmr.mgh.harvard.edu/) using each individual participant’s anatomical MRI scan. Initial MEG analyses were then performed in *Brainstorm* (Tadel, Baillet, Mosher, Pantazis, & Leahy, 2011). First the MEG sensor array was aligned with the anatomical model of the participant’s head using an automated error minimisation procedure. Covariance matrices were estimated from the data, and a head model comprising overlapping spheres was generated. A minimum norm solution was used to calculate a source model, with dipole orientations constrained to be orthogonal to the cortical surface. The model consisted of a set of linear weights at each location on the cortical surface that transformed the sensor space representation into source space.

MEG data were then imported into Matlab using *Fieldtrip* (Oostenveld, Fries, Maris, & Schoffelen, 2011), bandpass filtered (using the same filter as for the EEG data) and epoched. Univariate and multivariate analyses were performed in the same way as described for the EEG data in section 2.5. This was done using the sensor space representation (with 239 working sensors), the source space representation at approximately 500 vertices evenly spaced across the cortical mesh, and also within discrete regions of cortex defined by the Mindboggle atlas (Klein et al., 2017). For this latter analysis, the mean number of vertices in each cortical region is given in Table A1 in the Appendix. We conducted further analyses using multiple time-points as observations, at a single spatial (sensor or cortical) location.

## 3 Results

### 3.1 Experiment 1: EEG reveals above-chance classification of percepts

Mean event-related potentials (ERPs), averaged over the ten occipital electrodes where the changes in response from baseline were greatest (Figure 1a), showed a typical response to brief visual stimulation (black curve, Figure 1a). Clear ERPs were evident for all individual participants (thin traces, Figure 1a). In the grand average (black curve), two successive positive responses were evident over occipital electrodes at early time-points (126ms and 225ms after stimulus onset), corresponding to stimulus onset and offset. A later time-point (594ms after stimulus onset) showed negative voltages in occipital areas and positive voltages in frontal electrodes (see upper scalp plots for voltage distributions).

**Figure 1:**
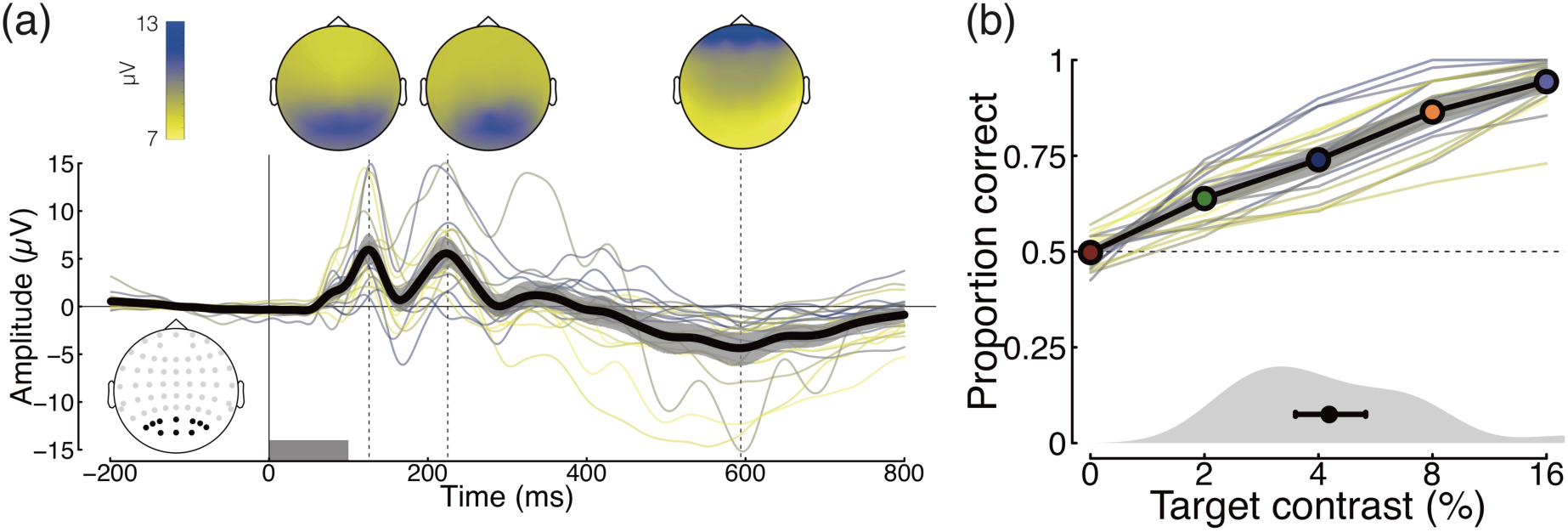
Grand mean ERPs (a) and summary of psychophysical performance (b). The black trace in panel (a) shows the grand mean across all conditions and participants (N=22, with 2000 ERPs per participant), with the grey shaded region giving 95% confidence intervals derived from 10000 bootstrap resamples. Thinner coloured traces show results for individual participants. In all cases the evoked responses were averaged across the 10 posterior electrodes shown in the lower left inset. The grey rectangle along the lower axis indicates the period during which the stimulus was presented. Scalp distributions of voltages at three time points (126ms, 225ms and 594ms, marked by dashed vertical lines) are shown at the top of the plot. The black line and coloured symbols in panel (b) show the mean psychophysical performance in each condition, averaged across participants (N=22), with the grey shaded region giving 95% confidence intervals derived by bootstrapping. Thinner coloured traces show results for individual participants, and symbol colour corresponds to those used to indicate target contrast conditions in subsequent figures. The grey curve at the foot shows the distribution of individual thresholds at the 75% correct point, with the black circle giving the mean, and error bars giving 95% confidence intervals.

Task performance in the five target contrast conditions ranged from chance in the 0% target contrast condition (where there was no correct answer as the ‘target’ interval was determined arbitrarily) to close to ceiling in the 16% target contrast condition (94% correct). Average data (black line) and results for individual participants (thin traces) are shown in Figure 1b, where it is evident that increasing target contrast improved performance for all participants. We fitted cumulative Gaussian functions to each participant’s data to estimate threshold contrast at 75% correct. The mean threshold was 4.25%, with the distribution shown at the lower axis of Figure 1b.

We first grouped ERP data according to contrast, and compared evoked responses in the null (pedestal only) and target (pedestal + target) intervals. The upper row of Figure 2a shows the ERPs averaged across occipital electrodes, with the null interval responses shown in black, and the target interval responses in colour. The middle row of Figure 2a shows the differences between these two ERPs, with horizontal lines at y=−1.5 indicating time points showing cluster-corrected significant differences. For a target contrast of 0%, the two stimuli are identical, and there are no meaningful differences between the waveforms (the two brief periods of significance are type I errors by definition). As target contrast increases, significant differences emerge between 100ms and 700ms post stimulus onset. These likely reflect both differences in early evoked responses, and also later decision-related components. Multivariate analyses across all 64 electrodes showed significant decoding only at the highest two target contrast levels (lower row of Figure 2a) within the same time window.

**Figure 2:**
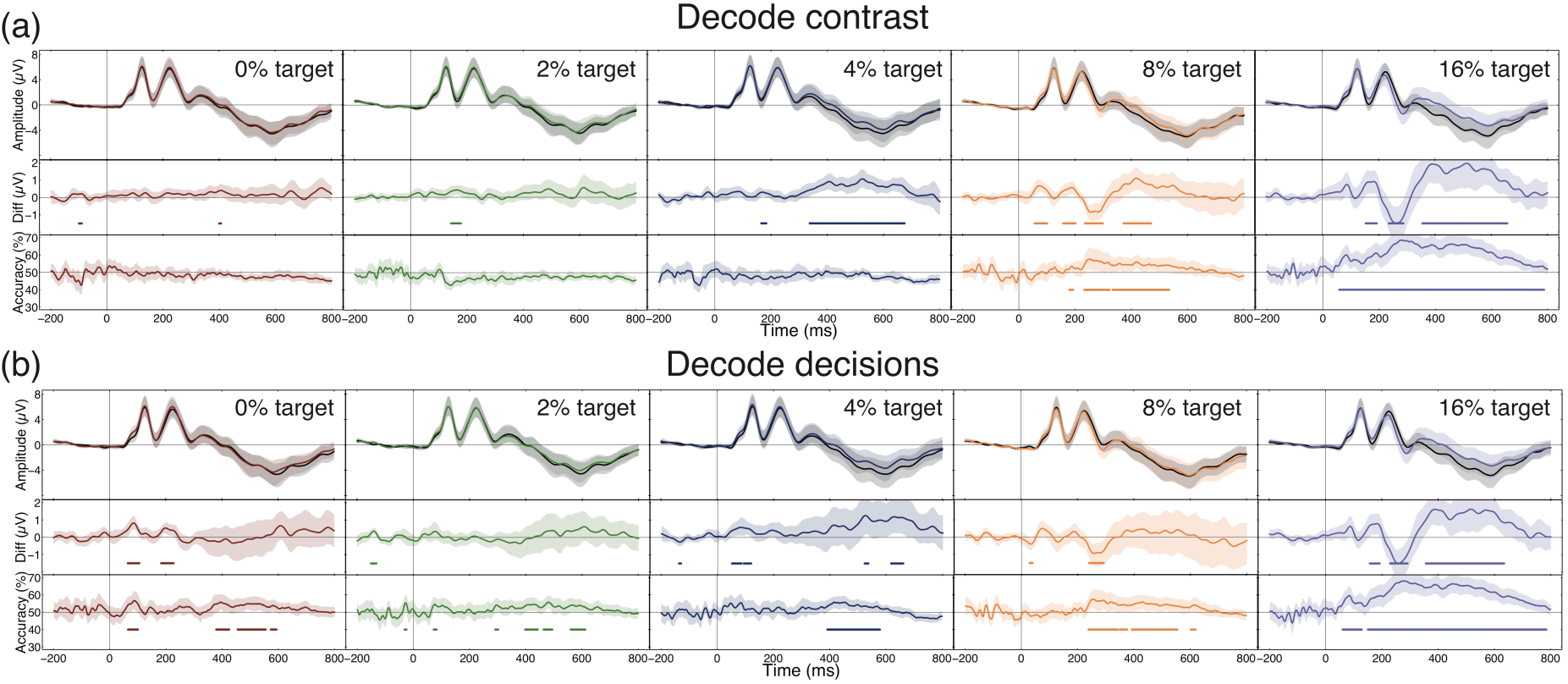
Univariate and multivariate analyses of EEG data. Panel (a) shows results for data partitioned according to the stimulus contrast (pedestal vs pedestal+target), and panel (b) shows results for data partitioned according to the participants’ perceptual decisions (selected vs non-selected). The upper section of each sub-plot contains grand averages of the ERPs being compared, in which the coloured curve indicates the target (or selected) waveform, and the black curve indicates the pedestal (or non-selected) waveform. The middle section of each sub-plot is the difference waveform. The lower section of each sub-plot shows multivariate classifier performance at each timepoint, where the baseline is 50% correct. In each panel, shaded regions show 95% confidence intervals across participants (N=22), calculated by bootstrapping. The coloured horizontal lines in the lower two sections indicate periods of time when the difference waveforms were significantly different from 0 (middle plots) or accuracy exceeded 50% correct (lower plots), calculated using a nonparametric cluster correction procedure (Maris & Oostenveld, 2007).

Next, we repeated the analyses on the same data, but this time organised according to the participant’s decisions rather than the physical stimulus contrast. In other words, we took ERPs from the intervals selected by the participants as appearing higher in contrast, and compared these with ERPs from the non-selected intervals. This analysis revealed additional time periods where the ERPs were significantly different, particularly in the 0% target condition, where differences were observed at around 100ms post stimulus onset. This finding was echoed in the multivariate analyses, which showed above chance decoding at early time points (around 100ms), as well as a sustained period of above chance decoding at all target contrasts from around 400-600ms post stimulus onset. The 0% target condition is of particular interest for this analysis, as any differences between evoked responses are not determined by the stimulus (which is identical in both intervals), and must be a consequence of differences in neural activity. The early significant clusters in both univariate and multivariate analyses indicate differences in the amplitude of the evoked response that influence subsequent perceptual decisions. Higher target contrasts increasingly converge with the contrast decoding analysis, as performance approaches ceiling (see Figure 1b) and the majority of selected intervals also contained the target (e.g. results for the 16% target condition are near identical in Figure 2a,b).

We tested the generality of the multivariate results in two ways. First, we took the classifier trained to discriminate between perceptual decisions at the highest target contrast (16%), and used this model to predict performance at lower target contrasts. This analysis (shown in Figure 3a) replicates the early periods of above chance decoding for 0% target contrast trials, suggesting that observers use a similar decision strategy for very challenging discriminations as for easier ones. Next, we took the classifier trained at each time point, and used it to predict selected and non-selected trials at all other time points (King & Dehaene, 2014). The results of this temporal generalization analysis (shown in Figure 3b,c) reveal isolated early structures around 100ms, and a more sustained pattern from 400-600ms (in the 0% condition) and from 200-800ms (in the 16% condition). We propose (see Discussion) that the early periods of above chance decoding may represent neural noise at the initial stages of processing, and the later periods could reflect noise in perceptual decisions, or memory traces from the first temporal interval.

**Figure 3:**
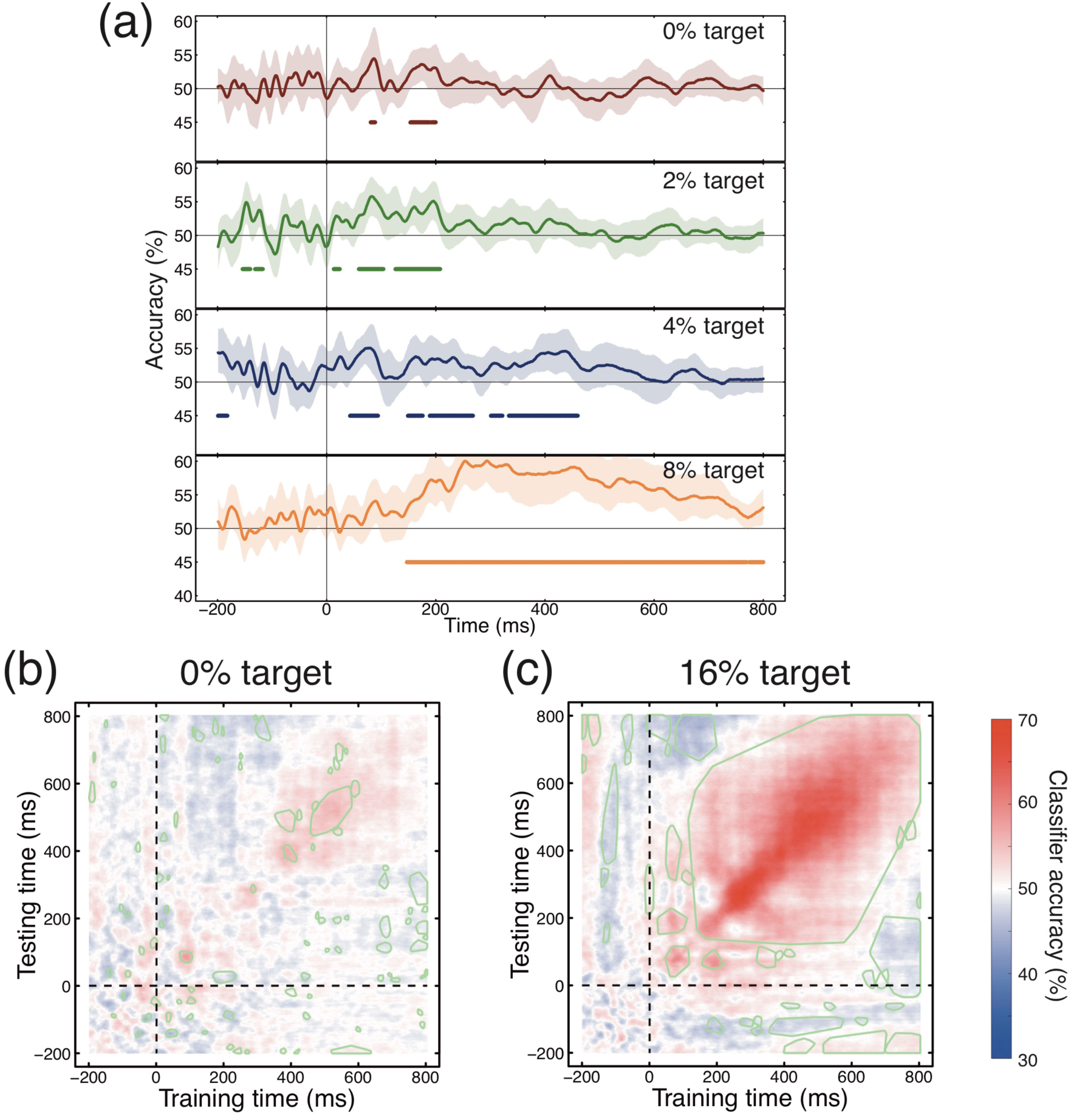
Multivariate generalization analyses across contrast condition (a) and time (b,c), for data partitioned according to the participants’ perceptual decisions. Panel (a) shows classifier accuracy at the four lower target contrasts after training the algorithm at the highest target contrast. Plotting conventions are as described for the multivariate analyses shown in Figure 2. Panels (b,c) show classifier accuracy when trained at each time point independently, and then tested using data at all time points. Regions outside of clusters where classifier accuracy differed significantly from chance (50% correct) are shaded.

### 3.2 Interval biases and decoding within the first or second interval

The temporal structure of a 2IFC trial is necessarily asymmetric, as the observer has knowledge of the first interval by the time they experience the second interval. In addition, repetition suppression effects can affect the evoked amplitude of the second presentation (Grill-Spector, Henson, & Martin, 2006). We first compared the average ERPs for all pedestal-only presentations (where the stimulus contrast was 50%) across the two intervals. We find both subtle and gross differences between these waveforms (see Figure 4a). Before stimulus onset, the waveforms differ as the second interval (green trace) has a decreasing voltage during the 200ms before the stimulus is presented. This likely originates from the tail end of the evoked response from the first interval (see Woldorff, 1993), which is decreasing from 400-600ms (the time window in which the second interval occurred). The second interval then has a more generally negative voltage throughout the 800ms following stimulus onset. The magnitude of this difference is much greater than that at stimulus onset, and so would persist even with a different baseline normalization regime (e.g. if the voltages were normalized to those at *t*=0). Furthermore, the differences become much more substantial at later time points, from 400-800ms. This may relate to the perceptual decision and motor response that the participant must make following the second interval.

**Figure 4:**
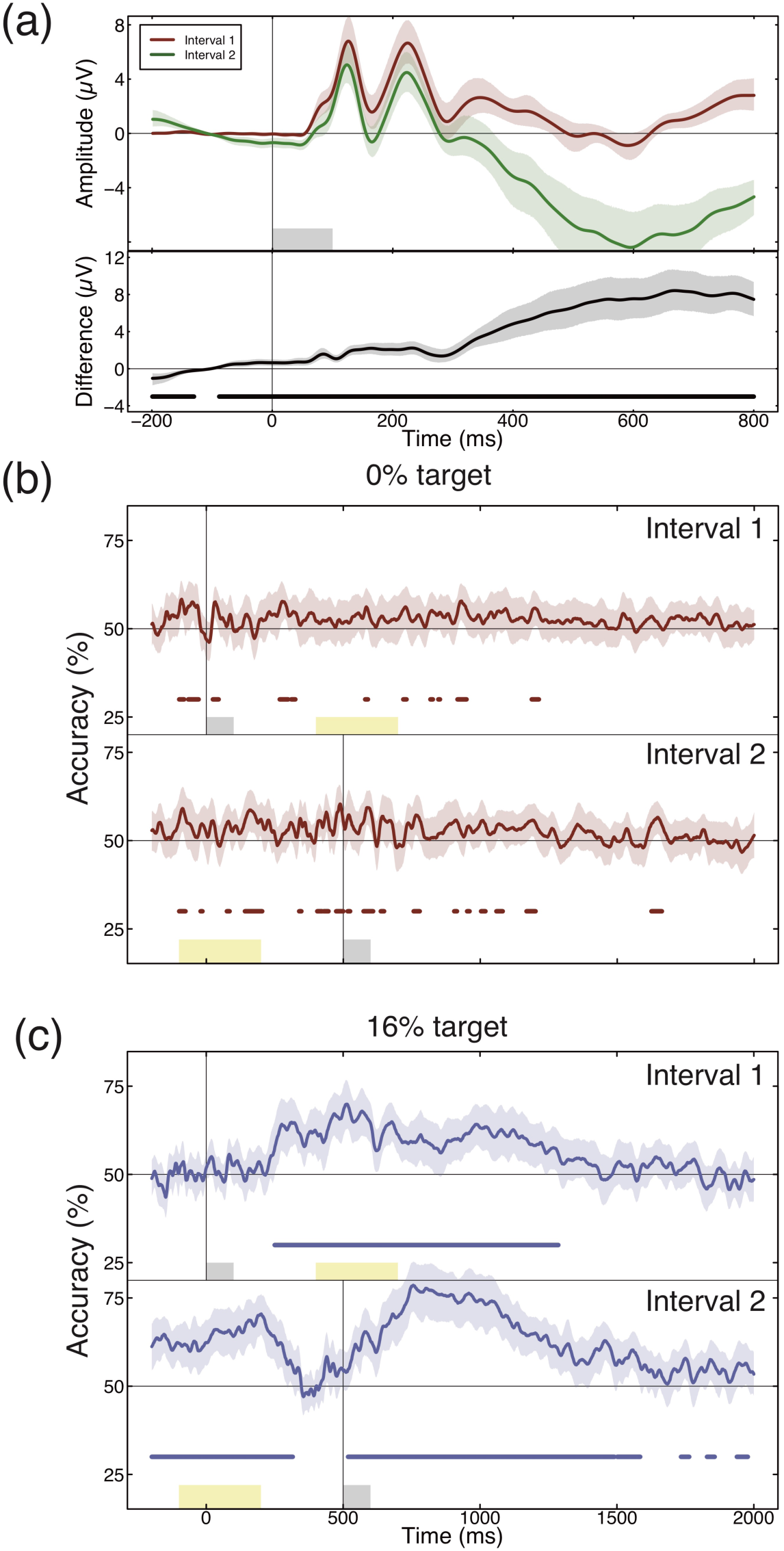
Comparisons across trial intervals. Panel (a) shows evoked responses in the first (red curve) and second (green curve) intervals of a 2IFC trial (upper plot) and their difference (lower plot). The evoked response is generally more negative in the second interval, particularly at time points >250ms after stimulus onset, despite the contrasts being physically identical (both 50%). Panels (b,c) show multivariate pattern classifier accuracy when comparing evoked potentials time-locked to either the first interval (upper plots) or the second interval (lower plots), for target contrasts of 0% (panel b) or 16% (panel c). In each plot, the shaded grey region shows the presentation of the stimulus from that interval and the yellow shaded region shows the range of time points when the stimulus from the other interval was presented (the precise inter-stimulus interval was jittered on each trial to reduce entrainment of ERP averages). In all panels shaded regions around each curve show 95% confidence intervals across participants, and horizontal coloured lines indicate significant clusters, consistent with conventions in previous figures.

Do these substantial differences in the evoked response to two physically identical stimuli affect the observer’s perception of the stimulus, or their decision over which interval to choose? We estimated interval bias for all participants by calculating the proportion of trials on which the second interval was selected, for the 200 trials in the 0% target contrast condition (where the two stimuli are identical). If this index is significantly below 0.5, it indicates a bias towards the first interval, and if it is significantly above 0.5 it indicates a bias towards the second interval. Despite individuals showing idiosyncratic biases (indices ranged from 0.23 to 0.92), the mean bias index was precisely 0.5 (SD: 0.14) and not significantly different from it (*t*_21_=0.11, *p*=0.91). The substantial voltage differences (Figure 4a) therefore do not appear to reflect group level differences in the appearance of the stimuli across intervals, and any idiosyncratic biases would presumably only reduce the power of our decision-based decoding analyses (Figure 2b), which are nevertheless significant.

The size of the voltage differences across intervals prompted us to investigate the extent to which decisions can be decoded within one or other interval, making comparisons across trials (within an interval) instead of across intervals (within a trial). The finite number of trials, combined with the presence of interval biases for some observers (see above) meant that there were often different numbers of trials available in the two intervals, so it was necessary to train and test the classifier on averages of fewer than 40 trials in some cases. The results of this multivariate analysis are shown in Figure 4b/c for decoding perceptual decisions at 0% contrast (Fig 4b) and at 16% contrast (Fig 4c), and for all conditions in Figure A1. In each sub-plot, the upper trace shows the classifier performance for data from interval 1 (with ERPs aligned at *t*=0ms), and the lower trace shows the classifier performance for data from interval 2 (with ERPs aligned at *t*=500ms). The yellow shaded regions indicate the time window when the stimulus in the other interval was displayed (the jittered inter-stimulus interval means that this time window is probabilistic rather than exact). Overall, we find increased decoding accuracy in the second interval compared with the first. This presumably reflects the increased information available for making a decision following the second stimulus.

### 3.3 Experiment 2: source space decoding is more sensitive than sensor space decoding

We confirmed that our MEG data replicated the key effects from Experiment 1 in several ways. First, we performed univariate and multivariate analyses in sensor space, using a cluster of 8 left-posterior sensors for the univariate analysis (defined as sensors located in the posterior portion of the helmet where activity at 110*ms* was significantly greater than 0), and all working sensors (N=239) for the multivariate analysis. The results of this analysis are shown in Figure 5a,c for the data split by participants’ perceptual decisions. Consistent with the EEG results, we find above chance pattern classification at early time points (∼100ms) as well as later >200ms. Second, we performed complementary analyses in MEG source space, using ERPs from pericalcarine cortex (corresponding to early visual cortex) for the univariate analyses, and a subset of 500 vertices across the entire cortical surface for the multivariate analyses. The results of this analysis are shown in Figure 5b,d. The general pattern of results is consistent with the sensor space analysis, though the shape of the ERP waveforms from pericalcarine cortex is somewhat different from those recorded in sensor space, with the peak of the onset and offset response appearing more prominent. Interestingly, we found that the multivariate analysis produced greater classification accuracy in source space (maximum of 80% correct) versus sensor space (maximum of 72% correct). We discuss possible reasons for this in the Discussion. Having confirmed that the multivariate source space analysis can decode perceived contrast, we next asked which brain regions contained information relevant to the task.

**Figure 5:**
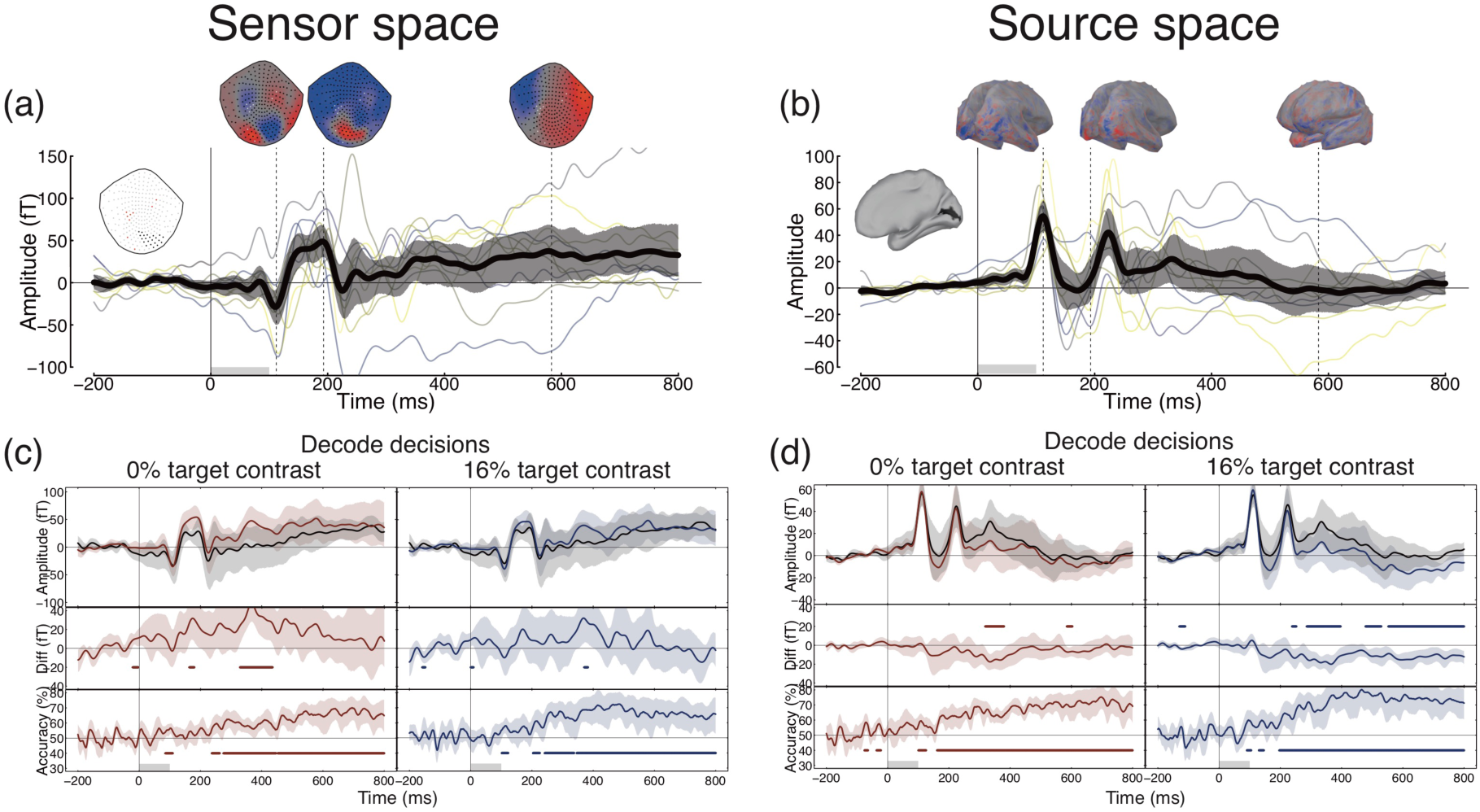
Sensor space and source space MEG analysis. Panel (a) shows the grand average ERP for all conditions and participants, pooled across subset of MEG sensors highlighted in black in black in the leftmost inset (red points are faulty sensors). Magnetic field distributions across the sensor array are shown at three time points at the top of the plot. Panel (b) shows a similar analysis in source space, for a region of cortex around the calcarine sulcus (highlighted black in the leftmost inset). The evoked response at each vertex on the cortical mesh was normalised such that the 110ms deflection was always positive, to avoid signal cancellation due to polarity inversions. In both panels, thin coloured curves represent individual participants (N=10). Panels (c,d) show univariate and multivariate comparisons between selected and non-selected ERPs in both contrast conditions, in the same format as described for Figure 2. Panel (c) shows this analysis in sensor space, and panel (d) shows the same analysis in source space. The source space multivariate analyses used a matrix of around 500 points distributed across the surface of the cortex.

### 3.4 Classification in anatomically-defined brain regions

We divided the cortex into 31 discrete non-overlapping anatomical regions using the *Mindboggle* atlas (Klein et al., 2017). (see Figure 6b). Maximal evoked potentials in these regions showed clear differentiation (see Figure A2). Because regions differed in size, each area contributed a different number of vertices on the cortical mesh for pattern classification (see Table A1).

At early time points, around 100ms, information in three adjacent regions around the occipital pole (the peri-calcarine region, the cuneus and the lateral occipital cortex) could be used to decode the participant’s percept in the 0% target contrast condition (final three traces in Figure 6a). Over time, this information spread forward to frontal and temporal cortex (see Figure 6c). By 300ms following stimulus onset, almost the entire brain contains information relevant to the task. This includes regions that do not appear to respond directly to presentation of visual stimuli (i.e. where there is no obvious evoked response, see Figure A2). A similar pattern of results is evident in the 16% target contrast condition (see Figure A3), confirming our earlier finding that differences in physical and perceived contrast are processed in a similar fashion.

**Figure 6:**
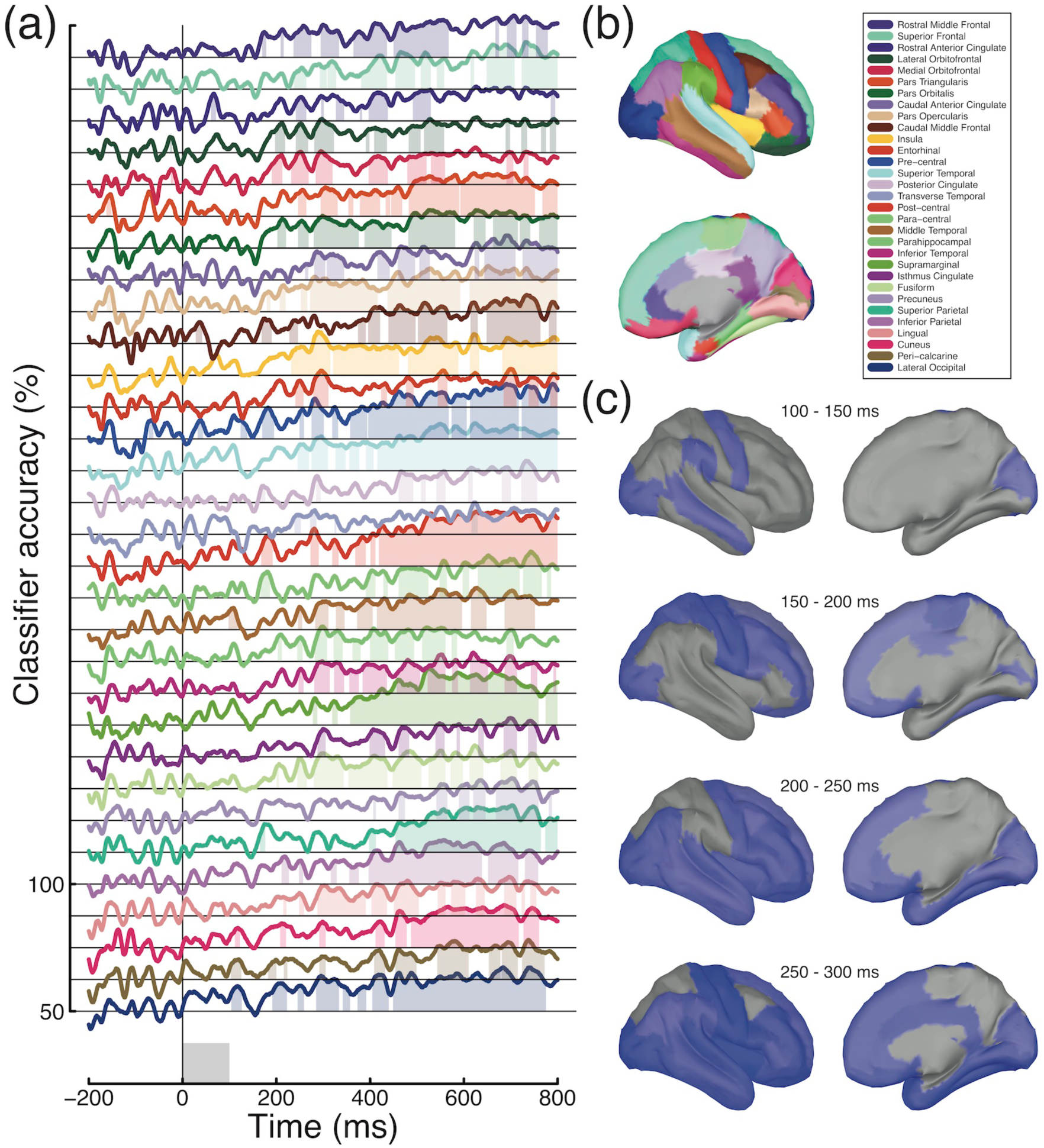
Atlas-based classification of decisions in the 0% target condition. Timecourses in panel (a) indicate classifier performance for each brain region, organised from anterior (top) to posterior (bottom) (see legend in panel b). These are offset vertically by 12.5% for each subsequent region for clarity, with the 50 and 100% labels relating to the bottom region but providing a common scale for each region. Shaded regions in panel (a) indicate clusters in which classification performance was significantly above chance (Bonferroni corrected for 31 brain regions). In panel (c), regions containing significant clusters within a given time window are shown in blue.

## 4 Discussion

The present study investigated the timecourse and location of perceptually relevant neural noise in contrast discrimination, using univariate and multivariate analysis of EEG and MEG data. Our results show that perceptual decisions are partly determined by responses in early visual cortex even when the two stimuli in a discrimination task are physically identical. This indicates that perceptually relevant neural noise impacts at the initial stages of processing and affects stimulus encoding in the visual system. However the best classifier performance occurred at later time points (>400ms), suggesting that additional sources of noise might also be involved. Analysis of differences across trial intervals revealed that neural activity in the second interval was more closely associated with subsequent decisions. We will now discuss the implications of these finding for our understanding of how neural activity (both evoked and spontaneous) influences the perceptual decisions involved in sensory discrimination.

### 4.1 Superior classification in MEG source space

Classifier performance overall was much higher for MEG data than for EEG data in identical conditions, despite the larger sample size of the EEG study (N=22 for EEG vs. N=10 for MEG). This is presumably due to the greater intrinsic sensitivity of MEG sensors, and the greater sampling density across the scalp (N=64 for EEG vs. N=239 for MEG). Classifier accuracy was also consistently higher in source space than in the sensor space representation primarily used in previous MEG studies (Cichy, Pantazis, & Oliva, 2014; Clarke, Devereux, Randall, & Tyler, 2015; Mostert, Kok, & de Lange, 2016). Since the source space representation is a weighted linear combination of activity at the sensors, this might be somewhat surprising. However, the source reconstruction presumably weights out signals from outside the brain (e.g. heart rate, breathing and blinking artefacts, and noise from outside of the scanner), resulting in a cleaner signal. Some form of source localisation may therefore be a useful processing step in future studies attempting multivariate classification of MEG signals. Additionally, combining the source space representation with atlas-based multivariate analysis permits questions to be asked about the information contained in specific brain regions at different points in time.

### 4.2 Single interval versus 2IFC

One distinction between this and most previous studies on the neural correlates of perceptual decision making is that previous work has used single interval (yes/no) paradigms (Hesselmann, Kell, Eger, & Kleinschmidt, 2008; Hillyard, Squires, Bauer, & Lindsay, 1971; Jolij, Meurs, & Haitel, 2011; Mostert et al., 2016; Ress & Heeger, 2003; Schölvinck, Friston, & Rees, 2012; Squires, Squires, & Hillyard, 1975), whereas here we used a 2IFC design. Since most psychophysical studies of contrast discrimination have used 2IFC, this choice has direct relevance to previous work. Additional benefits are that the number of evoked potentials in the selected and non-selected categories were necessarily balanced, and it was possible to analyse perceptual decisions based on two physically identical stimuli. In addition, 2IFC designs avoid problems with differences in bias (or response criteria) between participants, as pairs of stimuli are compared directly on a given trial (rather than against an internal standard). However, 2IFC cannot distinguish between hits and correct rejections (as these comprise ‘correct’ trials) or between misses and false alarms (incorrect trials), so direct comparisons of these trial categories is not possible in our design.

Another feature of 2IFC paradigms is that participants must hold information about the stimulus from the first interval in memory until after the second stimulus has been presented. This process may account for the sustained patterns of activity that permit classification long after stimulus offset (see Figures 2-6). In particular, our analysis of interval-specific effects (see Figure 4b,c) shows greater multivariate decoding accuracy in the second interval, presumably because at this point in the trial the observer has obtained all information necessary to make a decision.

### 4.3 Multiplicative noise

An alternative account of contrast discrimination performance at high pedestal contrasts is that transduction is linear but internal noise is signal-dependent (Pelli, 1985). If the dominant source of noise were early and multiplicative, this would avoid any issues relating to Birdsall’s theorem, as the transducer could be linear. It has proven difficult to distinguish between the multiplicative and additive noise accounts purely from contrast discrimination experiments (Georgeson & Meese, 2006; Kontsevich, Chen, & Tyler, 2002). At a single neuron level there is well-established evidence of multiplicative noise (Tolhurst, Movshon, & Dean, 1983), yet it appears that across populations of neurons with different sensitivities the overall noise is effectively additive (Chen, Geisler, & Seidemann, 2006). Since evidence from fMRI (Boynton et al., 1999), EEG (Busse et al., 2009) and psychophysics (Kingdom, 2016) all argue strongly against a linear transducer, we think this explanation is unlikely to account for the body of available data.

### 4.4 Resolving early noise and Birdsall’s theorem

Early noise has typically been considered at very early stages, including photoreceptor noise in the retina (Barlow, 1962), which can be considered as external noise (albeit in a different sense from experimentally added external noise, as it is not under the direct control of the experimenter). Late additive noise is often assumed (either implicitly or explicitly) to be added at the decision stage, long after the nonlinearities of early visual processing (Cabrera, Lu, & Dosher, 2015; Mueller & Weidemann, 2008). The results here point to a perceptually-relevant source of noise that is present in the early evoked response, at around 100ms or earlier. However we find that classification performance improves after this point in processing, reaching a maximum approximately 400-600ms after target onset (see Figures 2 % 5). In addition, our temporal generalisation analysis (see Figure 3b,c) shows that these two time windows involve distinct patterns of neural activity, implying separate sources of noise. This is consistent with a sequence of multiple (and presumably independent) noise sources at different stages of processing. Since mathematical treatment of complex systems involving multiple nonlinearities and noise sources is currently lacking, it is unclear what implications this would have for the visibility of early nonlinearities.

One possibility is that a strong source of noise occurs immediately after the initial contrast transduction nonlinearity in V1, leaving that nonlinearity visible but obscuring later ones. This would explain why psychophysical contrast perception maps closely onto the neural response from early visual areas (Baker & Wade, 2017; Barlow, Hawken, Parker, & Kaushal, 1987; Boynton et al., 1999), but not the highly compressive contrast-invariant response in later regions (Avidan et al., 2002; Rolls & Baylis, 1986). Indeed, this might enable the visual system to harness the properties of Birdsall linearisation to preserve the dynamic range of early representations through later processing (that is more compressive) when making comparisons across stimuli (as in a discrimination paradigm). Object recognition, and other operations that benefit from invariance to features such as contrast, position and size, but do not require comparisons across multiple stimuli, would be immune to the Birdsall effect and benefit from the later nonlinearities. Furthermore, a strong early source of noise would make the study of later ‘mid-level’ visual processes much more challenging, perhaps explaining why vision research has typically focussed on earlier mechanisms and can be caricatured as being ‘stuck’ in V1 (Graham, 2011; Peirce, 2007).

In order to investigate these possibilities further, we performed two additional analyses. To link the internal fluctuations measured in our experiments with a psychophysical measure of internal noise, we correlated classifier accuracy with the contrast discrimination thresholds estimated from the psychophysical responses in Experiment 1. Since high internal noise should result in higher discrimination thresholds (poorer performance), we predicted that the two measures would be correlated at time points where the neural fluctuations were most relevant to perception. This analysis is shown in Figure 7, and reveals a time window with significant negative correlations around 450-650ms (i.e. high thresholds correspond to poor classifier performance). We speculate that neural noise within this time window most closely corresponds to the ‘late’ additive noise that is a feature of contemporary models of contrast discrimination. However it is also possible that other factors mediate this relationship, including the interval bias described in the Results section which could inflate negative correlations by driving thresholds up and classifier performance down. Nevertheless, it is interesting to demonstrate a link between psychophysical thresholds and decoding of neural responses.

**Figure 7:**
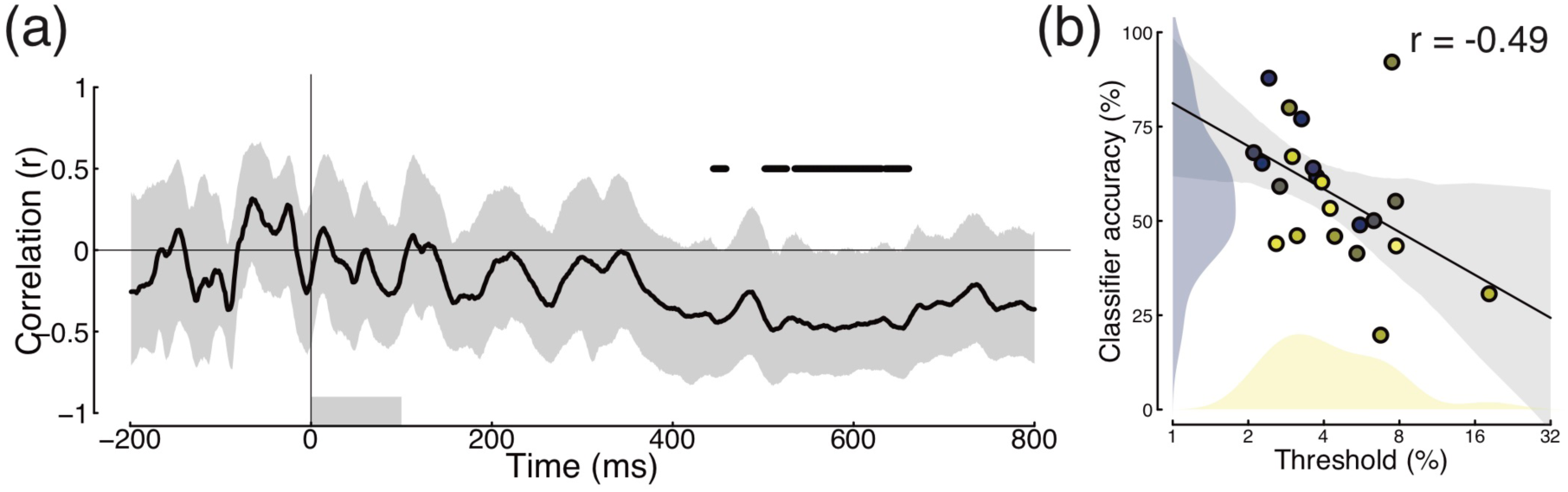
Correlation between individual contrast discrimination thresholds (see distribution in Figure 1b) and classifier accuracy in the 0% target contrast condition of Experiment 1 (N=22). Panel (a) shows the correlation as a function of time. The horizontal black lines at r=0.5 denote clusters of significant effects (two-tailed), according to a nonparametric cluster correction procedure (Maris & Oostenveld, 2007). Grey shaded regions represent bootstrapped 95% confidence intervals calculated across participants, and the lower grey rectangle shows the period when the stimulus was displayed. To further illustrate this relationship, panel (b) shows a scatterplot of the correlation between thresholds and the averaged classifier performance within all significant clusters identified in (a). The diagonal line is the best fitting Deming regression line, with grey shaded regions showing bootstrapped 95% confidence intervals, and blue and yellow histograms showing the distribution of values for each measure.

The final analysis we performed was inspired by the suggestions of an anonymous reviewer, who pointed out that in our main multivariate analyses, although the classifier is always trained on information from both trial intervals, test data are supplied from one interval at a time. This means that the classifier’s decisions differ from those of human participants, who in a 2IFC paradigm always have information available from both trial intervals. We conducted further multivariate analyses, by training and testing the classifier on downsampled timecourses of entire 2IFC trials combined across both intervals (to account for the jittered ISI, each interval was aligned to its respective trigger).

The results of this analysis are shown in Figure 8 for the EEG experiment (Figure 8a,c), and for the MEG experiment in both sensor space (Figure 8b,d) and source space (Figure 8e,f). All data sets produced above-chance classification at some sensors and brain regions, indicating that patterns across time were able to discriminate neural states. For the 16% target contrast, early visual areas at the occipital pole showed high classifier accuracy (Figure 8f), consistent with the salient target contrast increment producing greater ERP amplitudes in the target interval (see Figures 2 & 5). For the 0% target contrast condition, accuracy in early visual regions was relatively poor, and the highest accuracy was in fronto-parietal regions (Figure 8e). This suggests that the most important signals for classifying decisions in this condition arise after the initial responses in visual brain areas. The later sustained response from around 400ms onwards (see Figures 2b & 3b) seems more consistent with the brain regions producing significant decoding here. These additional analyses suggest that the internal noise sources most relevant for contrast discrimination performance occur subsequent to the initial visual cortical responses, and are therefore more consistent with models of ‘late’ noise than with early internal (or unintended external) noise.

**Figure 8:**
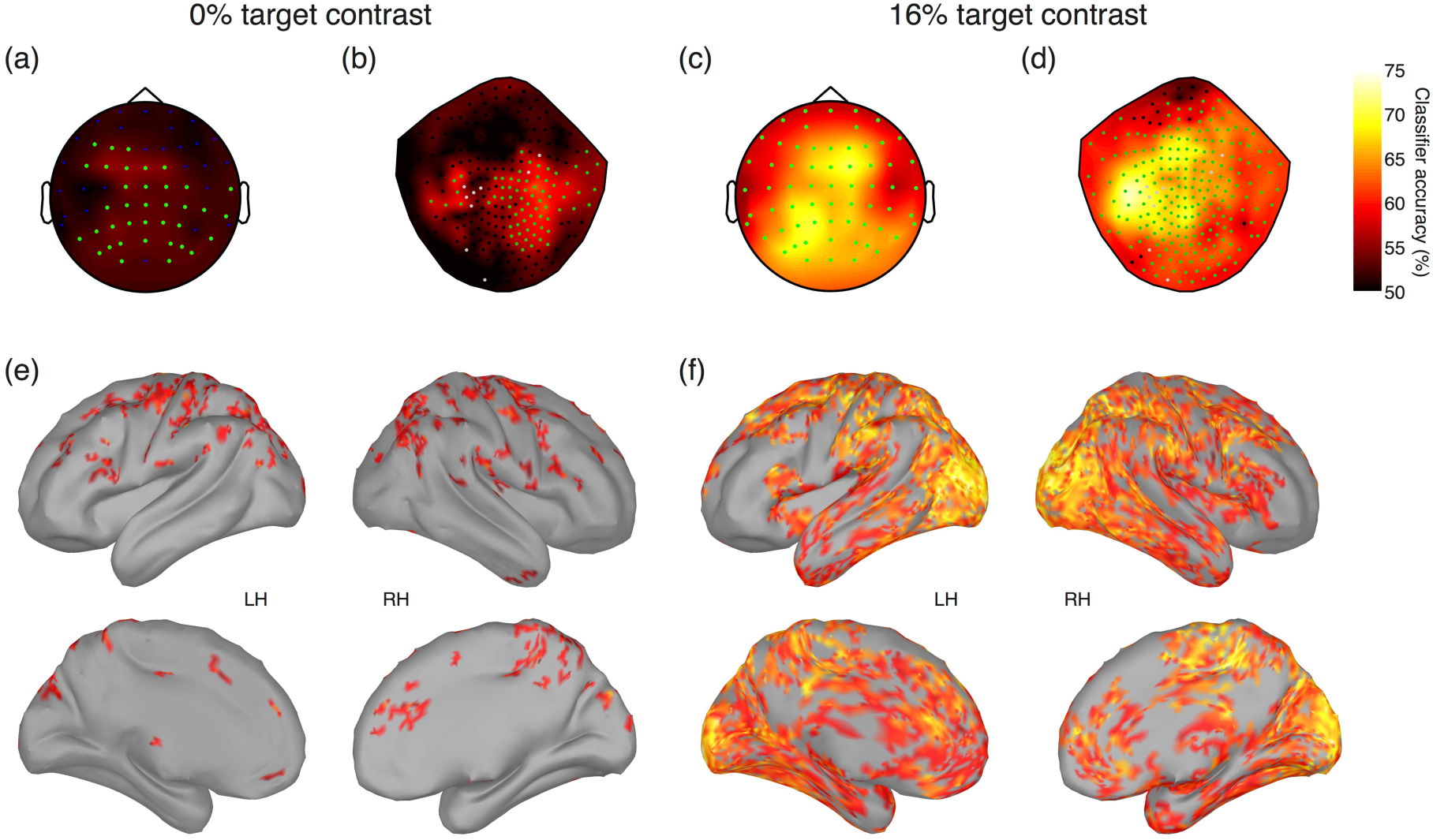
Whole-trial pattern classification accuracy in sensor space and source space. Entire time courses of both 2IFC intervals were categorised according to participant percepts (selected vs non-selected), and downsampled in steps of 10ms. Classifier accuracy was averaged across participants at individual electrodes in the EEG experiment (panels a,c), in sensor space in the MEG experiment (panels b,d) and at 15000 vertices on the cortical surface in source space in the MEG experiment (panels e,f). Sensors comprising significant clusters are marked by green points in panels a-d, and vertices not part of significant clusters are coloured grey in panels e,f.

### 4.5 Conclusion

To summarise, in this study we investigated the timecourse of the neural operations involved in contrast discrimination. We demonstrated that internal noise impacting early in time (around 100ms after stimulus onset) and in the visual pathway can affect sensory processing and perceptual decisions. However, the strongest internal noise source was later (around 400-700ms), involved parietal and frontal brain regions, and was correlated with psychophysical thresholds. Our novel application of multivariate analysis methods to discrete spatial regions of MEG source space offers the capability of studying how the brain represents information in both space and time.

### Data and code availability statement

Analysis scripts, experiment code and raw EEG data are publicly available on the *Open
Science Framework* website, at the following URL: http://doi.org/10.17605/OSF.IO/GBHQJ

MEG data are not publicly available for this project. The reason for this is that ethical approval and informed consent were not sought for sharing these data publicly. In particular the structural MRI scans could in principle be used to identify individuals (e.g. by generating a 3D model of their faces). It may be possible to share these data with individual researchers, subject to ethical approval from the York Neuroimaging Centre. Please contact Dr Daniel Baker if you wish to pursue this.

## 6 Appendices

**Figure A1:**
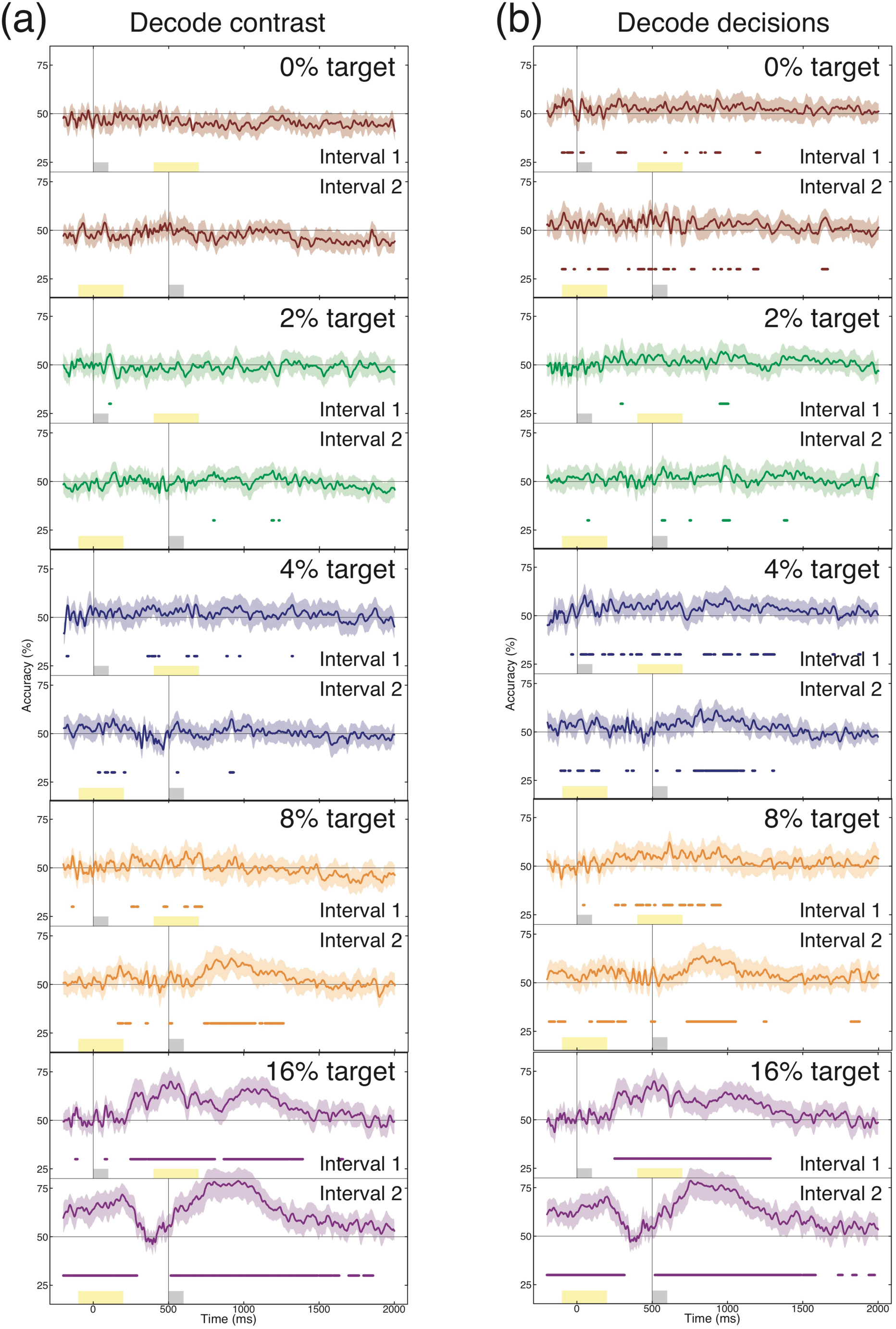
Interval-based MVPA analysis for all target contrast conditions, and for both contrast-based decoding (a) and decision-based decoding (b). Plotting conventions are consistent with Figure 4b,c.

**Figure A2:**
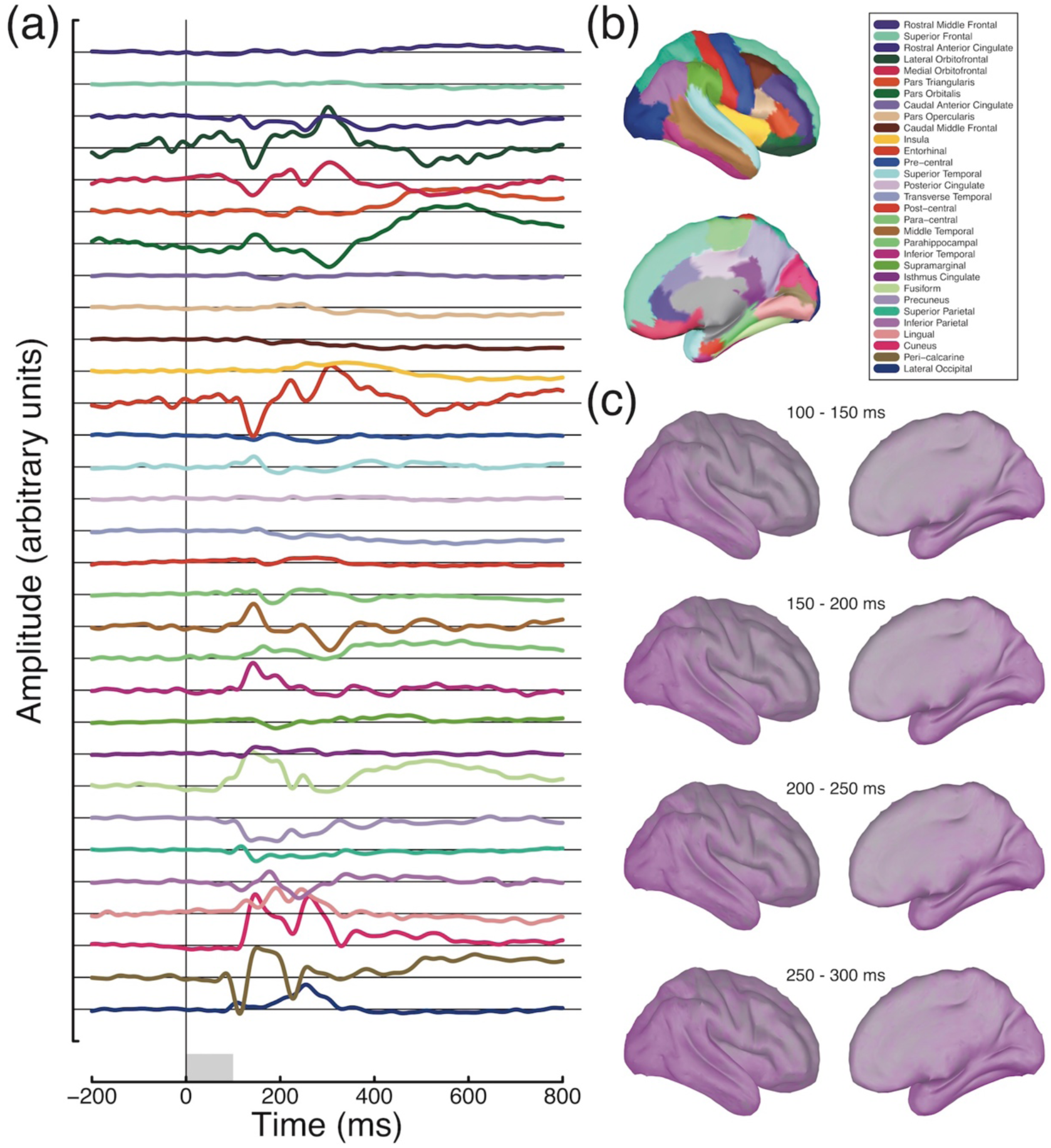
Maximal evoked responses in different anatomical regions. Each trace in panel (a) plots the timecourse of the vertex in the named region (see legend in panel (b)) with the largest absolute deflection from baseline. Panel (c) shows absolute activity averaged across four time windows, demonstrating that the majority of activity occurs in occipito-temporal regions.

**Figure A3:**
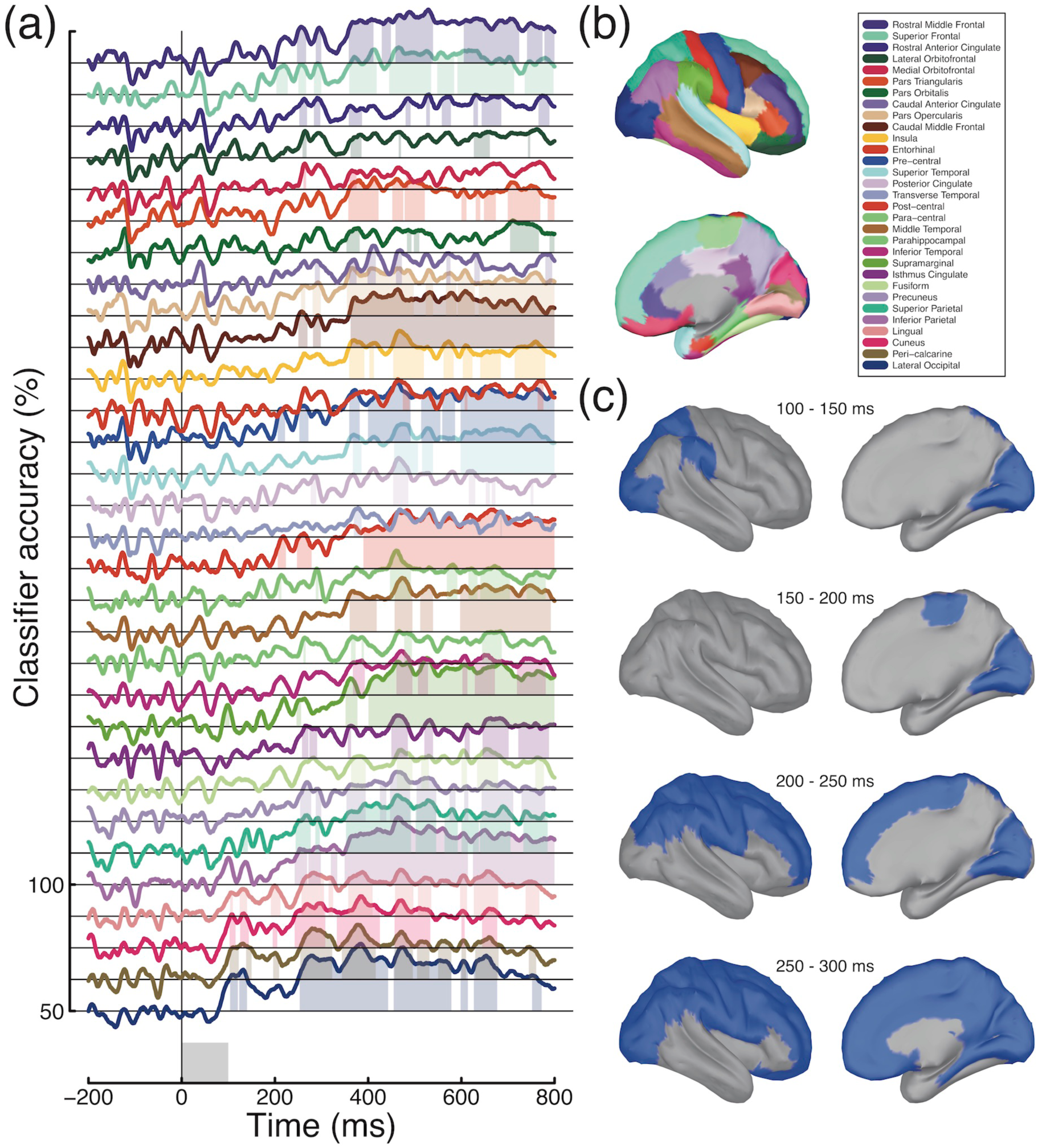
Atlas-based classification of decisions in the 16% target condition. Plotting conventions mirror those of Figure 6.

**Table A1:** Numbers of vertices on the cortical mesh. Individual regions were taken from a mesh consisting of around 3000 vertices, and pooled across hemispheres. The ‘whole brain’ mesh (final row) was subsampled to around 500 vertices. Precise numbers of vertices varied across individual participants owing to individual differences in brain size and morphology. Entries in the ‘Colour’ column correspond to the colours used in Figures 6, A2 & A3.

| Region | Colour | Mean size | Minimum size | Maximum size |
| --- | --- | --- | --- | --- |
| Rostral Middle Frontal |  | 157 | 145 | 173 |
| Superior Frontal |  | 342 | 317 | 384 |
| Rostral Anterior Cingulate |  | 31 | 27 | 37 |
| Lateral Orbitofrontal |  | 100 | 83 | 113 |
| Medial Orbitofrontal |  | 53 | 45 | 62 |
| Pars Triangularis |  | 63 | 56 | 69 |
| Pars Orbitalis |  | 31 | 27 | 35 |
| Caudal Anterior Cingulate |  | 29 | 25 | 34 |
| Pars Opercularis |  | 53 | 43 | 61 |
| Caudal Middle Frontal |  | 75 | 59 | 91 |
| Insula |  | 60 | 53 | 69 |
| Entorhinal |  | 16 | 9 | 25 |
| Pre-central |  | 140 | 124 | 155 |
| Superior Temporal |  | 166 | 150 | 183 |
| Posterior Cingulate |  | 38 | 32 | 45 |
| Transverse Temporal |  | 9 | 6 | 11 |
| Post-central |  | 142 | 130 | 156 |
| Para-central |  | 48 | 41 | 61 |
| Middle Temporal |  | 134 | 123 | 152 |
| Parahippocampal |  | 21 | 17 | 25 |
| Inferior Temporal |  | 118 | 96 | 149 |
| Supramarginal |  | 123 | 102 | 155 |
| Isthmus Cingulate |  | 28 | 23 | 34 |
| Fusiform |  | 81 | 73 | 87 |
| Precuneous |  | 119 | 93 | 140 |
| Superior Parietal |  | 148 | 136 | 168 |
| Inferior Parietal |  | 149 | 131 | 157 |
| Lingual |  | 101 | 69 | 125 |
| Cuneus |  | 67 | 58 | 74 |
| Peri-calcarine |  | 38 | 24 | 45 |
| Lateral Occipital |  | 162 | 139 | 181 |
| Whole brain |  | 503 | 503 | 504 |

